## Supplementary figures for "Bioprinting microporous functional living materials from protein-based core-shell microgels"

Author information

1 State Key Laboratory of Materials-oriented Chemical Engineering, College of Chemical Engineering, Nanjing Tech University, 30 Puzhu South Road, Nanjing, 211816, P.R. China.

2 Yusuf Hamied Department of Chemistry, University of Cambridge, Lensfield Road, Cambridge, CB2 1EW, UK.

3 Cambridge University-Nanjing Centre of Technology and Innovation, 126 Dingshan Street, Nanjing 210046, P. R. China.

4 State Key Laboratory of Materials-oriented Chemical Engineering, College of Biotechnology and Pharmaceutical Engineering, Nanjing Tech University, 30 Puzhu South Road, Nanjing, 211816, P.R. China.

5 Cancer Research UK Cambridge Institute, University of Cambridge, Cambridge, Li Ka Shing Centre, Robinson Way, Cambridge, CB2 0RE, UK

6 Cavendish Laboratory, University of Cambridge, J J Thomson Avenue, Cambridge, CB3 0HE, UK

7 Current address: Department of Chemical Engineering, University College London, Torrington Place, London, WC1E 7JE, UK

*

**Supplementary figures**

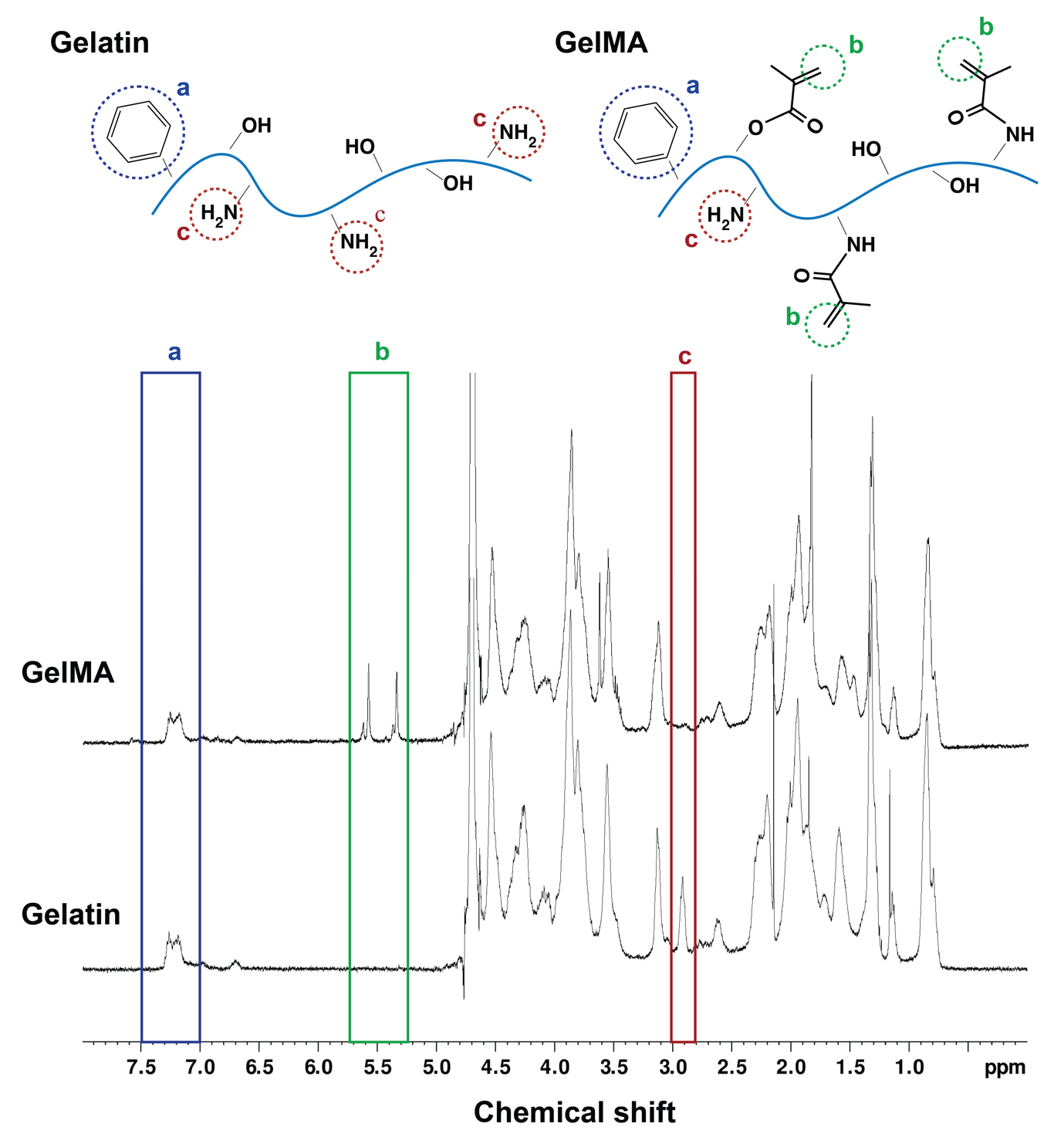

**Fig. 1. GelMA synthesis.** ^1^H NMR spectra of gelMA and gelatin. The substitution gives rise to a noticeable new proton signal from 5.4 t0 5.7 ppm (methacryloyl signal, b) whilst a reduction around 2.9 ppm (lysine signal, c). The degree of functionalization is estimated by normalizing lysine signals in both spectra by respective aromatic proton signals (7.0-7.5 ppm, a).

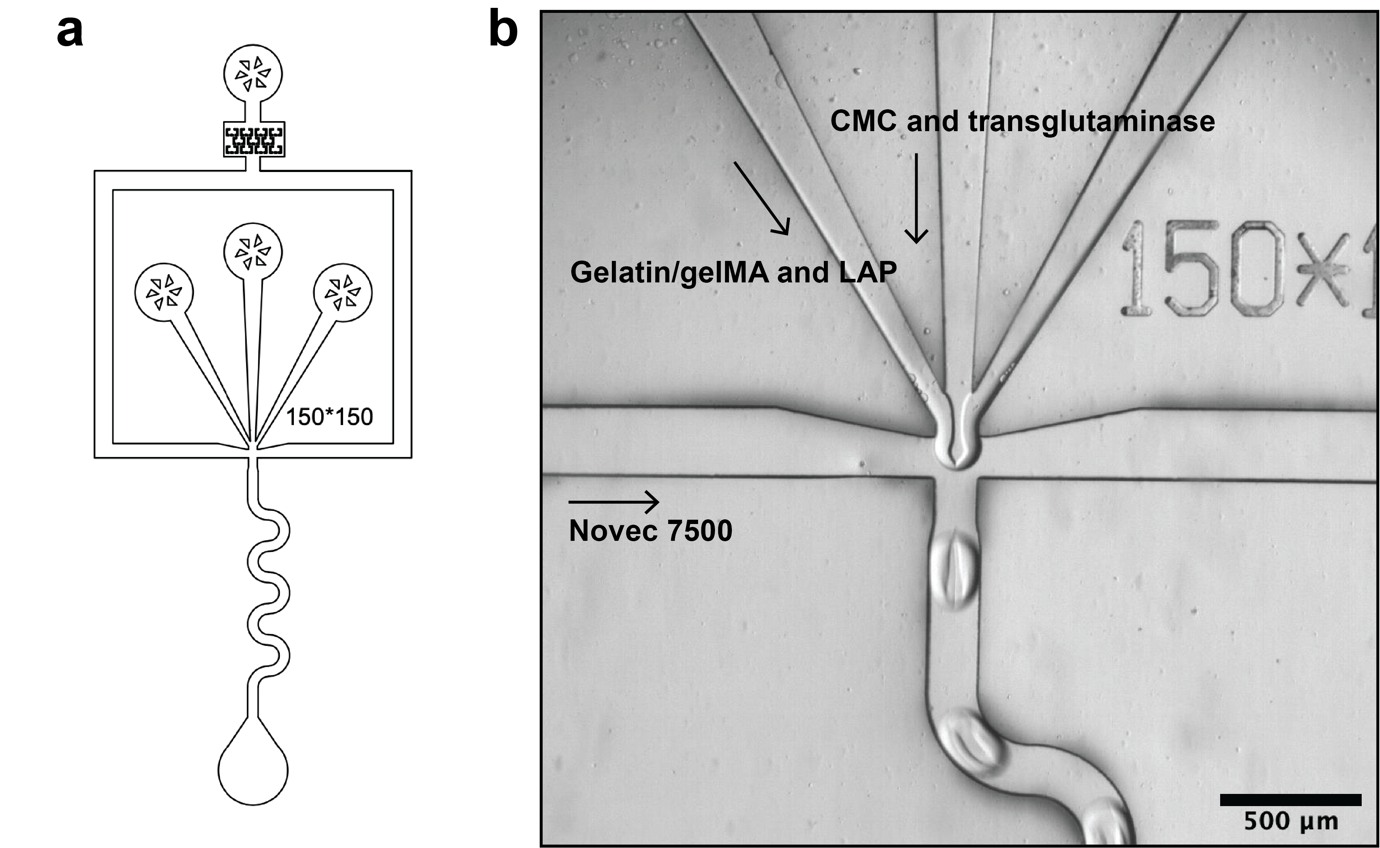

**Fig. 2. Droplet microfluidics.** (a) Design of the microfluidic device used for droplet generation. (b) Micrograph of the core-shell droplet generation.

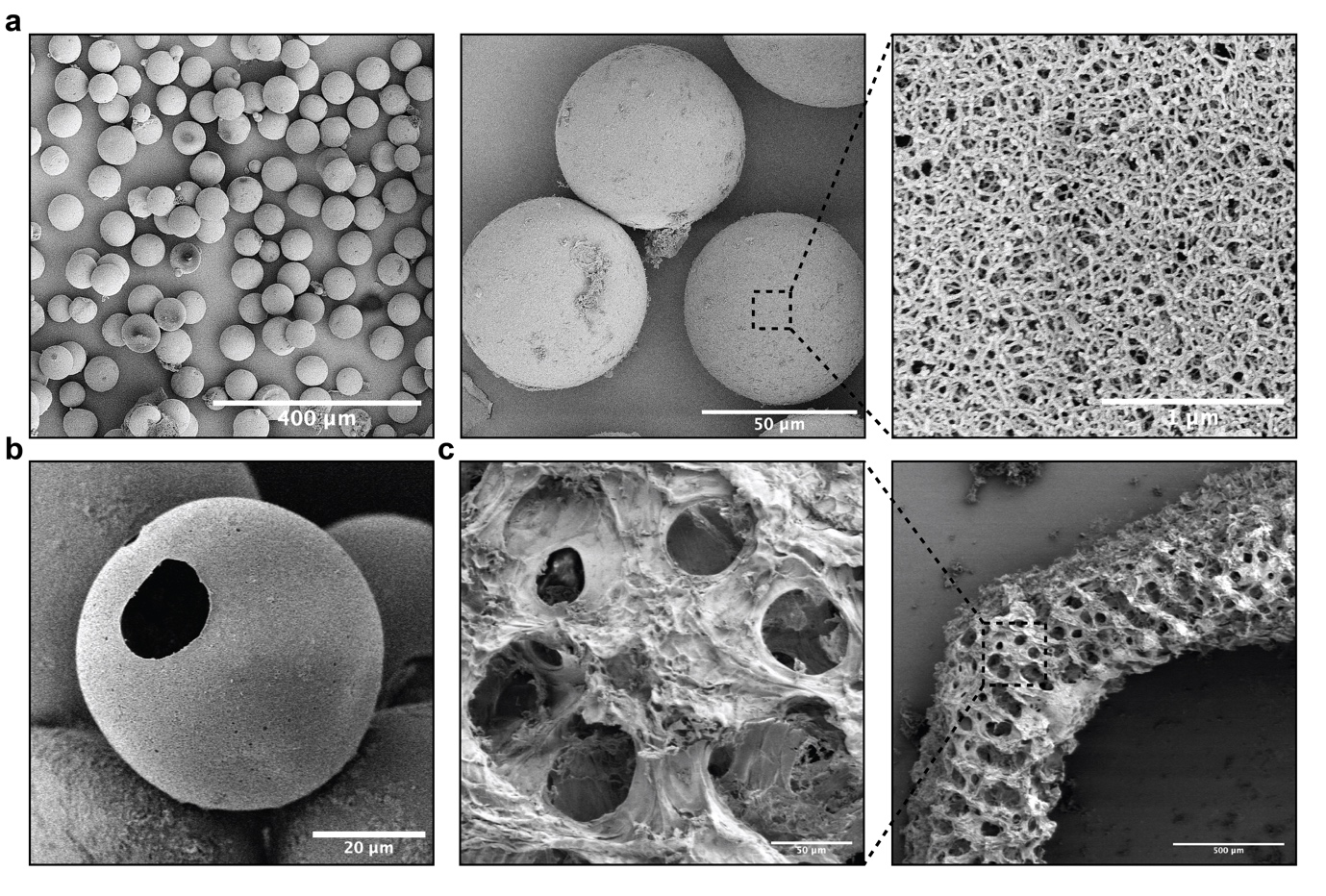

**Fig. 3. Scanning Electron Microscope.** (a)-(b) Surface morphology of microgels. Samples were prepared by critical point drying. (a) Nanoporosity of the microgel surface. (b) Dehydrated core-shell microgels with a solid shell and a hollow core. (c) Surface morphology of PAM scaffolds. Samples were prepared by lyophilization. During drying, microgels tended to shrink; however, due to the interconnectivity of microgels, the structure of single microgels were flattened and hence micropores were revealed.

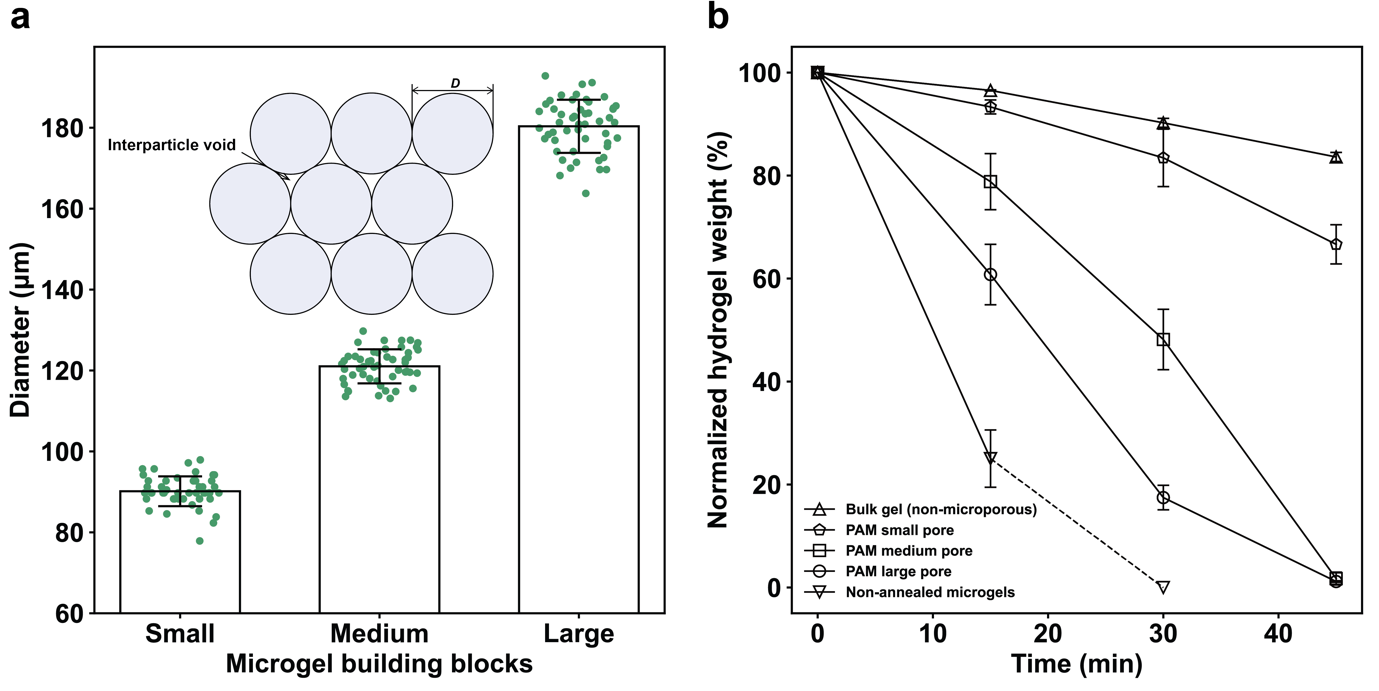

**Fig. 4. Scaffold degradation kinetics by percolation of trypsin.** (a) Size distribution of microgels of varying diameters. Small: 90.2±3.7 µm, medium: 121.0±4.21 µm, and large: 180.4±6.57 µm, *n* = 50 microgels. (b) Digestion of scaffolds assembled from differently sized microgels and of dual-crosslinked bulk hydrogels and freely suspended microgels. The non-annealed microgels are completely liquified in 30 min, *n* = 3 independent experiments. All data are presented as mean values +/- standard deviation.

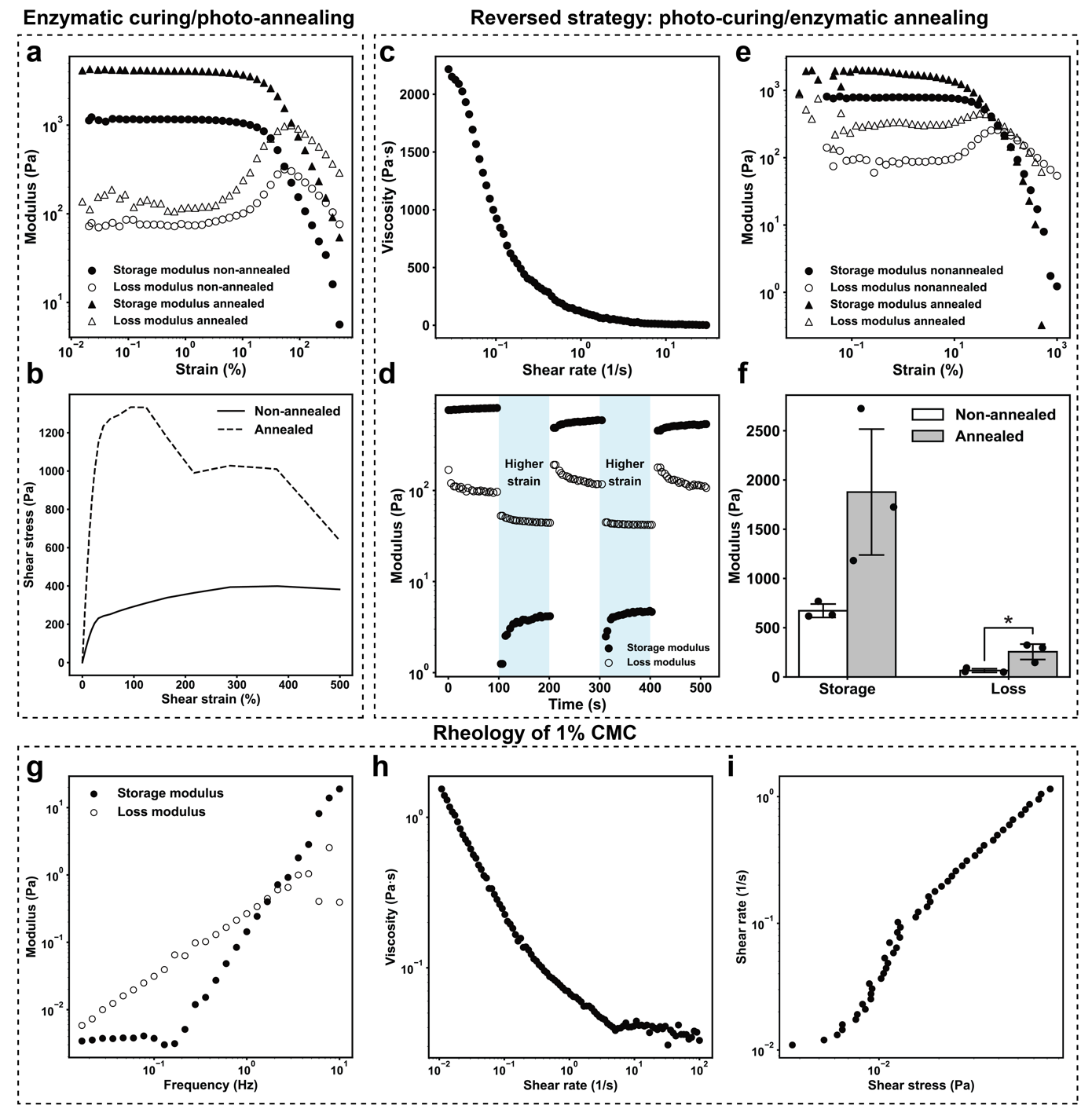

**Fig. 5. Rheology.** (a)-(b) Rheological characterization of jammed microgel inks and PAM scaffolds (enzymatic curing and photo-initiated annealing). (a) Strain sweep test. Strains from 0 to 1000% and frequency 1 Hz. (b) Stress-strain curve. (c)-(f) Rheological characterization of PAM scaffolds with a reversed crosslinking strategy (photo-initiated curing and enzymatic annealing). (c) Shear rate sweep experiment. Shear rate from 0 to 10 1/s. Strain 1%. (d) Step-strain sweep where low strain (1%) and high strain (90%) are cycled every 100 s. Frequency 1 Hz. (e) Strain sweep test. Strains from 0 to 1000% and frequency 1 Hz. (f) Storage and loss moduli before and after enzymatic annealing. Data are presented as mean values +/- standard deviation, *n* = 3 independent experiments. *p=0.028, unpaired two-tailed student’s t test. (g)-(i) Rheological characterization of 1% CMC. (g) Frequency sweep experiment. Frequency from 0 to 10 Hz. Strain 1%. (h) Shear rate sweep experiment. Shear rate from 0 to 100 1/s. Strain 1%. (i) Shear rate-shear stress curve.

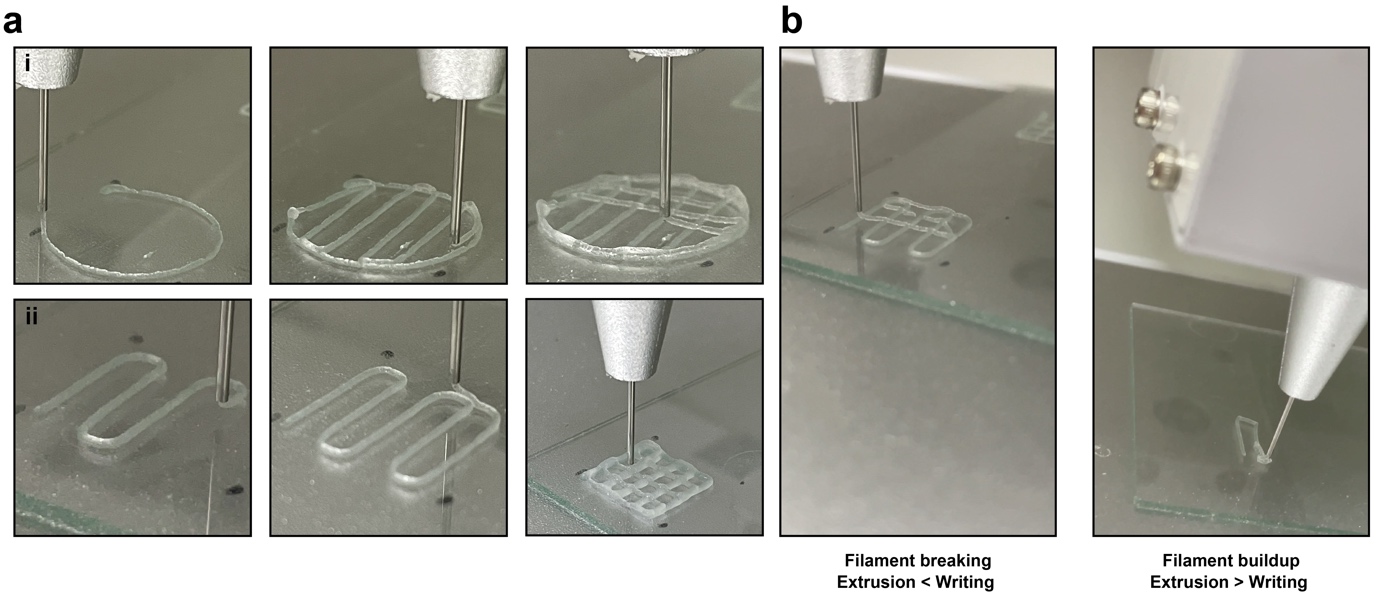

**Fig. 6. Extrusion printing of jammed microgels.** (a) Using a commercial 3D printer, jammed microgels could be patterned into predefined shapes, such as a (i) circle or a (ii) square. (b) Failure of printing owing to mismatched speed. When writing speed is markedly larger than extrusion, filament breaking happens, and when extrusion speed is larger than writing, the filament unfavorably builds up.

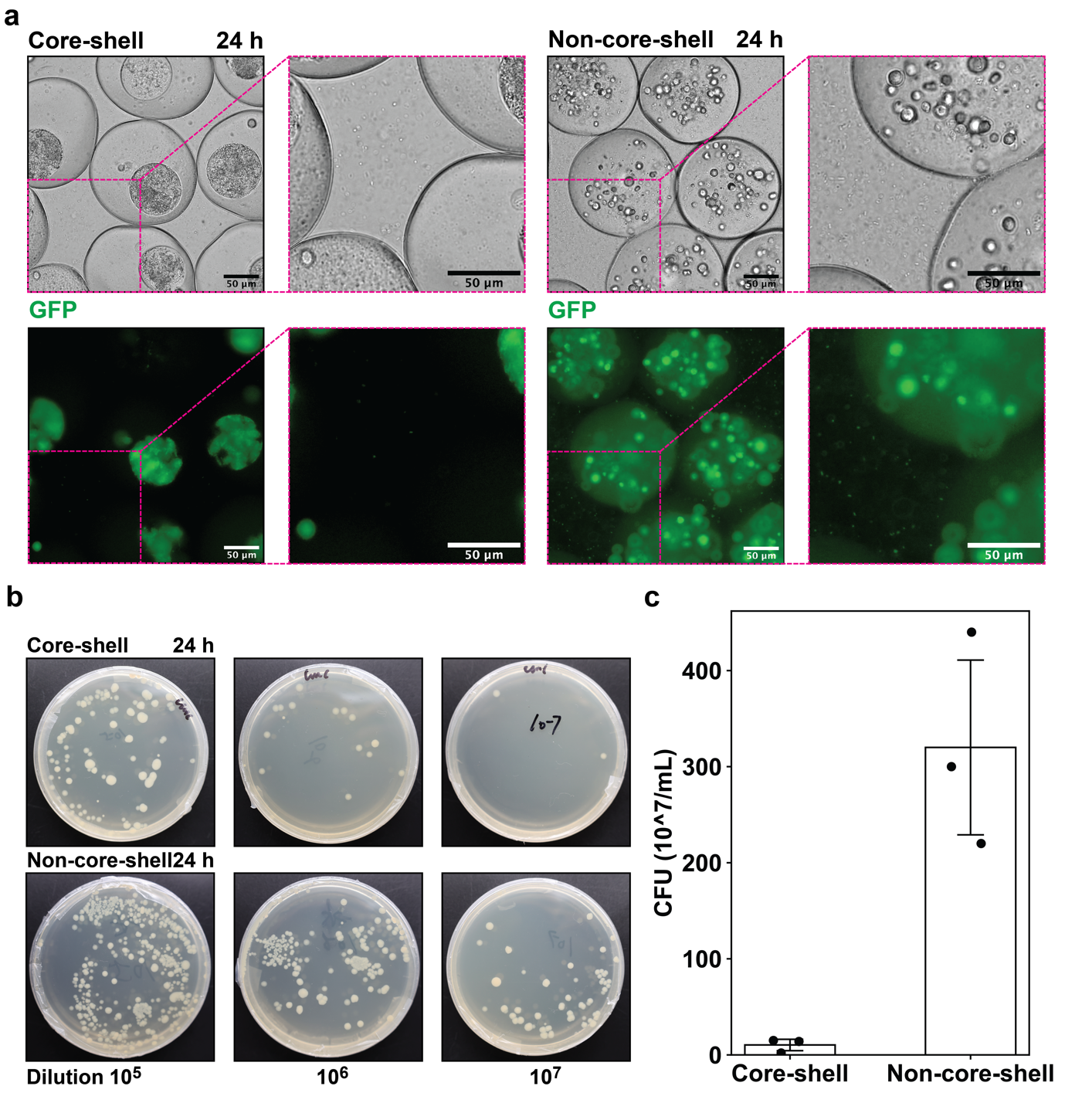

**Fig. 7. Core-shell microgels mitigate cell leakage.** (a) Local magnification of Fig. 3a and Fig. 3b in the main content. Higher level of *E. coli* leakage can be visually identified in the non-core-shell microgels. (b) The dilution plating experiment with core-shell and non-core-shell microgels. (c) Calculated CFU. Data are presented as mean values +/- standard deviation *n* = 3 independent biological experiments.

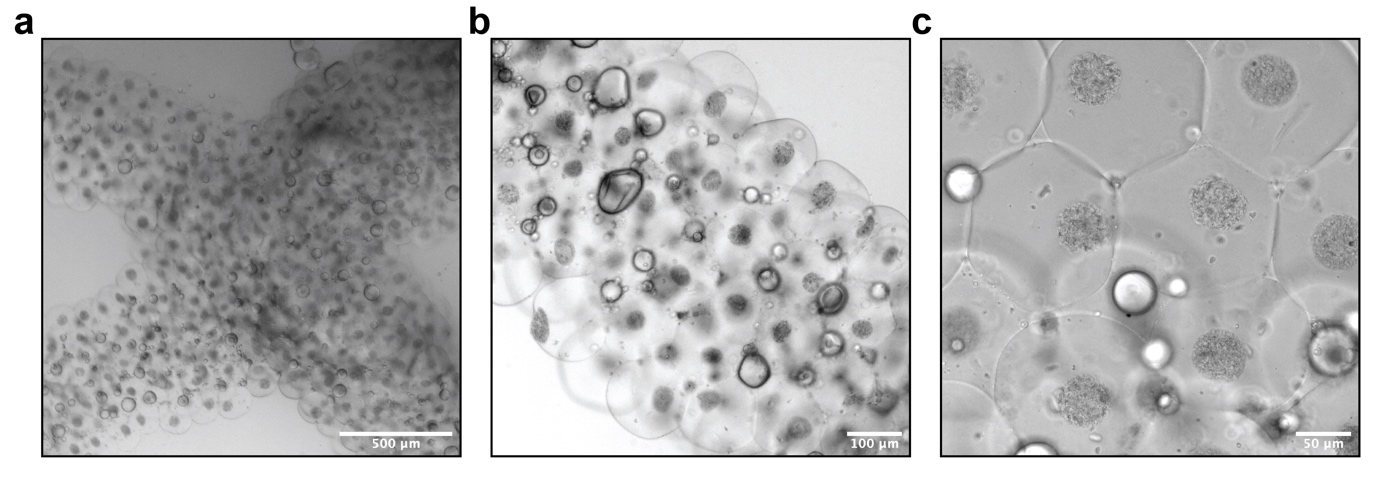

**Fig. 8. Scaffold for microbial culture.** The scaffold immobilizes the consortium of *Chlorella vulgaris* and *Bacillus subtilis* in separate microgels after 24 hours of cell culture in BG11 media. Also, as shown from (c), the micropores can be seen under a brightfield microscope after the swelling of PAM scaffolds.

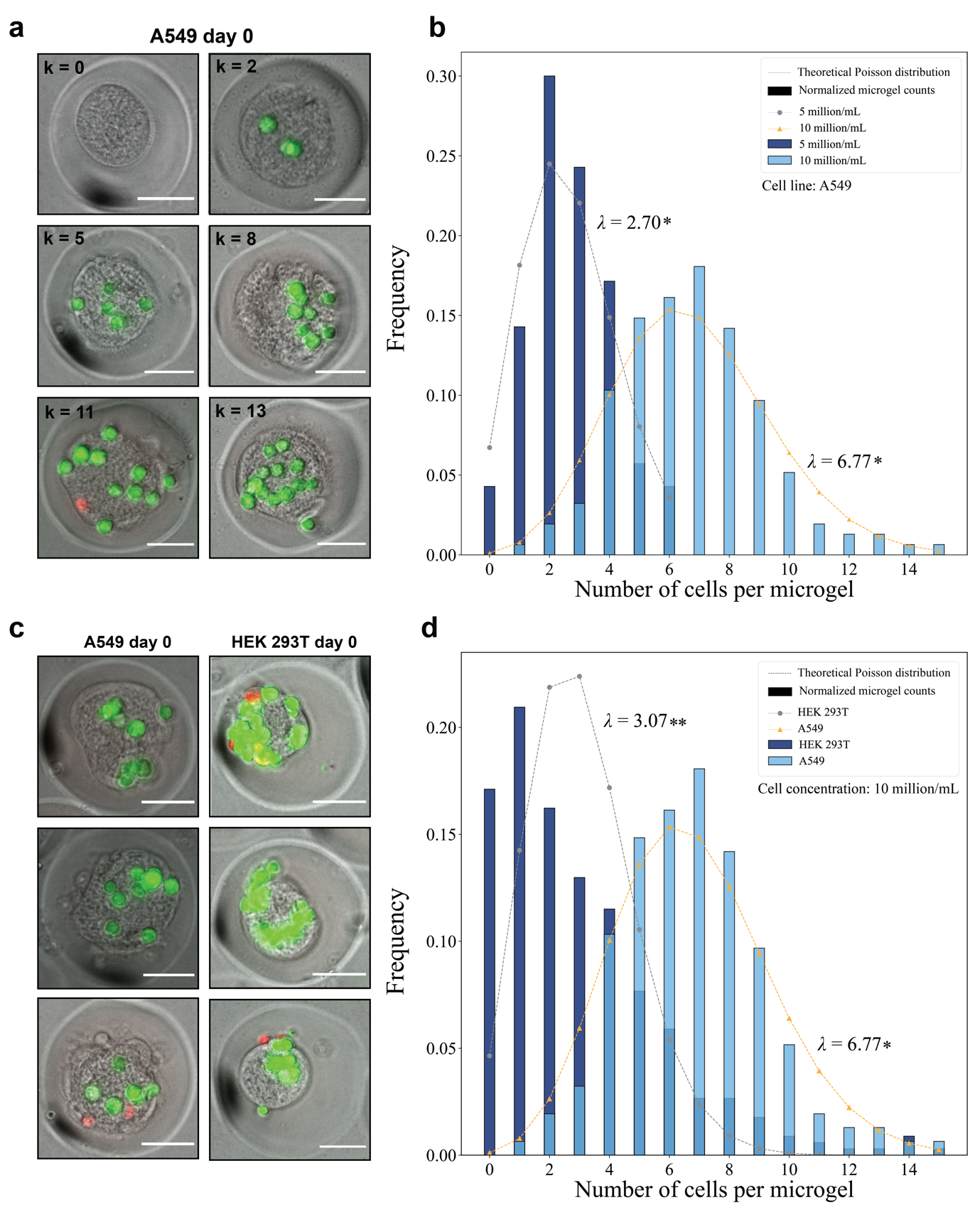

**Fig. 9. Poisson encapsulation.** (a) Micrographs showing the variety of cell numbers (A549 cells) inside microgels after cell encapsulation. *k* represents the number of cells encapsulated. (b) The observed frequency of A549 cell cumbers per microgel co-plotted with the theoretical Poisson distribution. The encapsulation of A549 follows the Poisson stochasticity, verified by two cell concentrations, namely, 5 and 10 million cells/mL. * p>0.05, chi-square test. (c) Comparison of A549 and HEK 293T encapsulation, which shows that the latter tends to aggregate to form cell clusters and hence disrupts the Poisson stochasticity. (d) The observed frequency of HEK 293T and A549 cell cumbers per microgel co-plotted with their theoretical Poisson distribution. Cell concentration is 10 million/mL, which shows that the encapsulation of HEK 293T cells does not follow a typical Poisson distribution. * p=0.999, and ** p=0.003, chi-square test. Scale bars in a and c represent 50 µm.

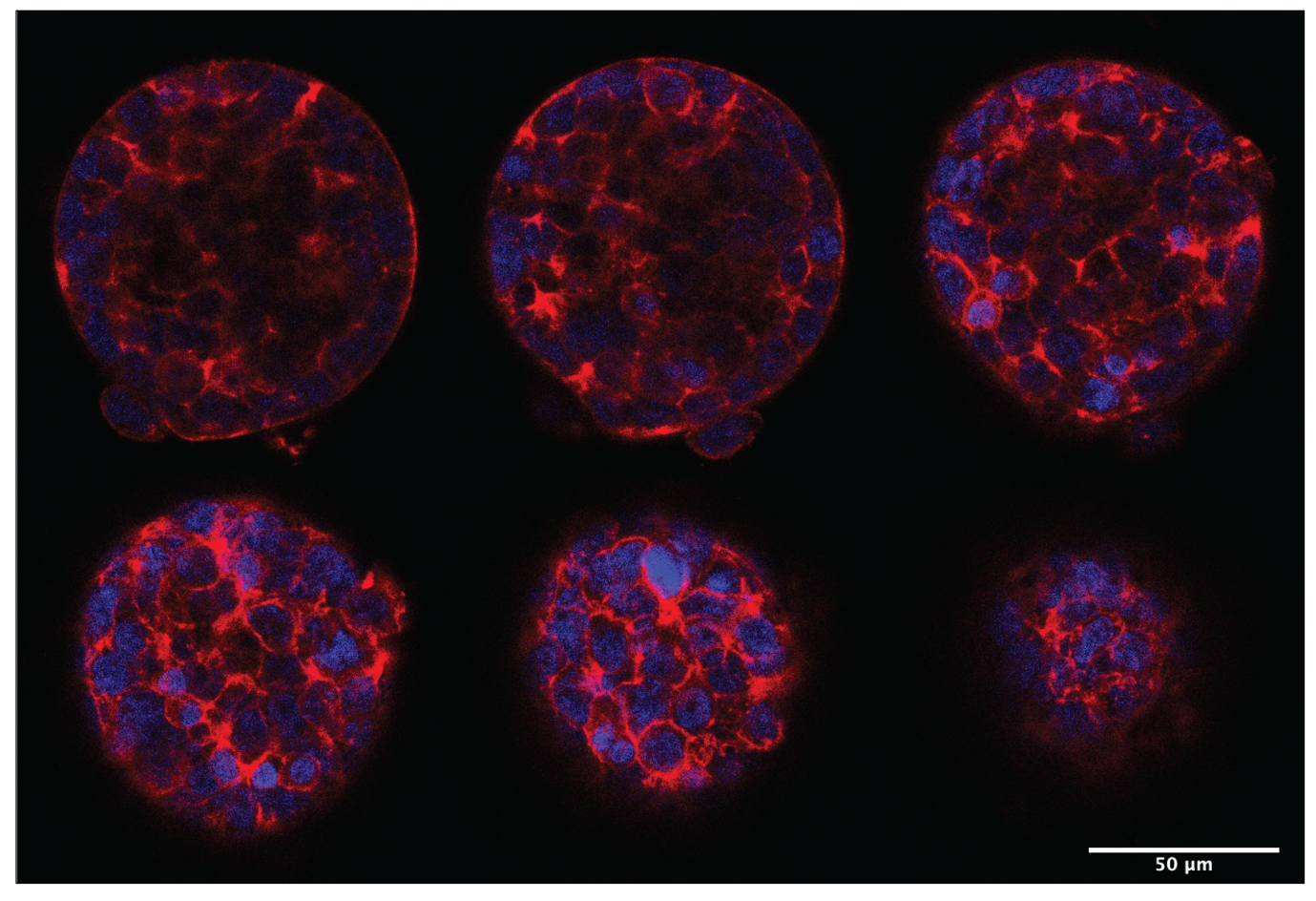

**Fig. 10. Cytoskeletal structure of HEK 293T spheroids.** Confocal microscopic images at different vertical locations of a HEK 293T spheroid stained with DAPI and phalloidin.

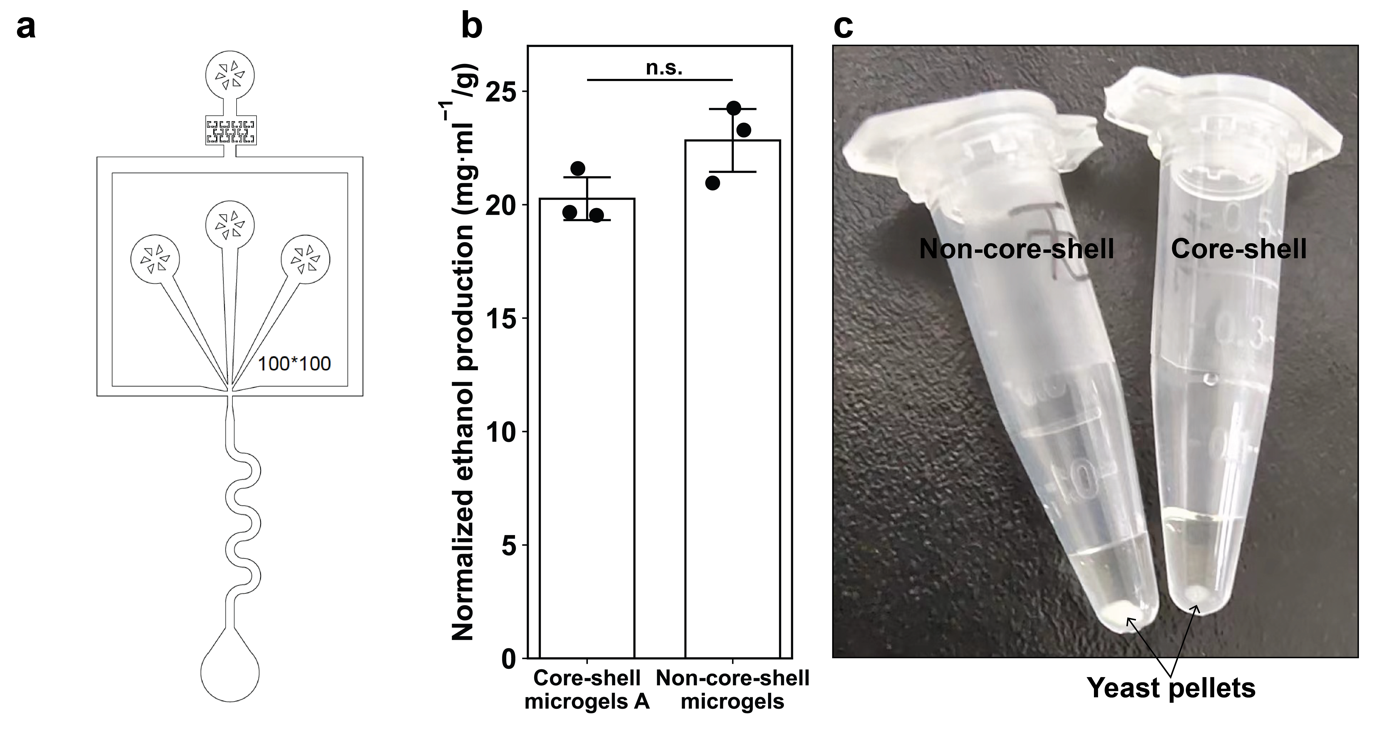

**Fig 11. Yeast-laden core-shell microgel scaffolds for ethanol fermentation.** (a) Design of the microfluidic used to generate smaller core-shell microgels C. (b) The normalized ethanol production at 20 hours. Data are presented as mean values +/- standard deviation, *n* = 3 independent experiments, unpaired two-tailed student *t* test. (c) Before gas chromatography, the sampled media (50 µL) were centrifuged to remove the yeast cells. It can be seen that the yeast pellet from the medium in which the core-shell microgel scaffold functioned is smaller than its core-shell counterpart.

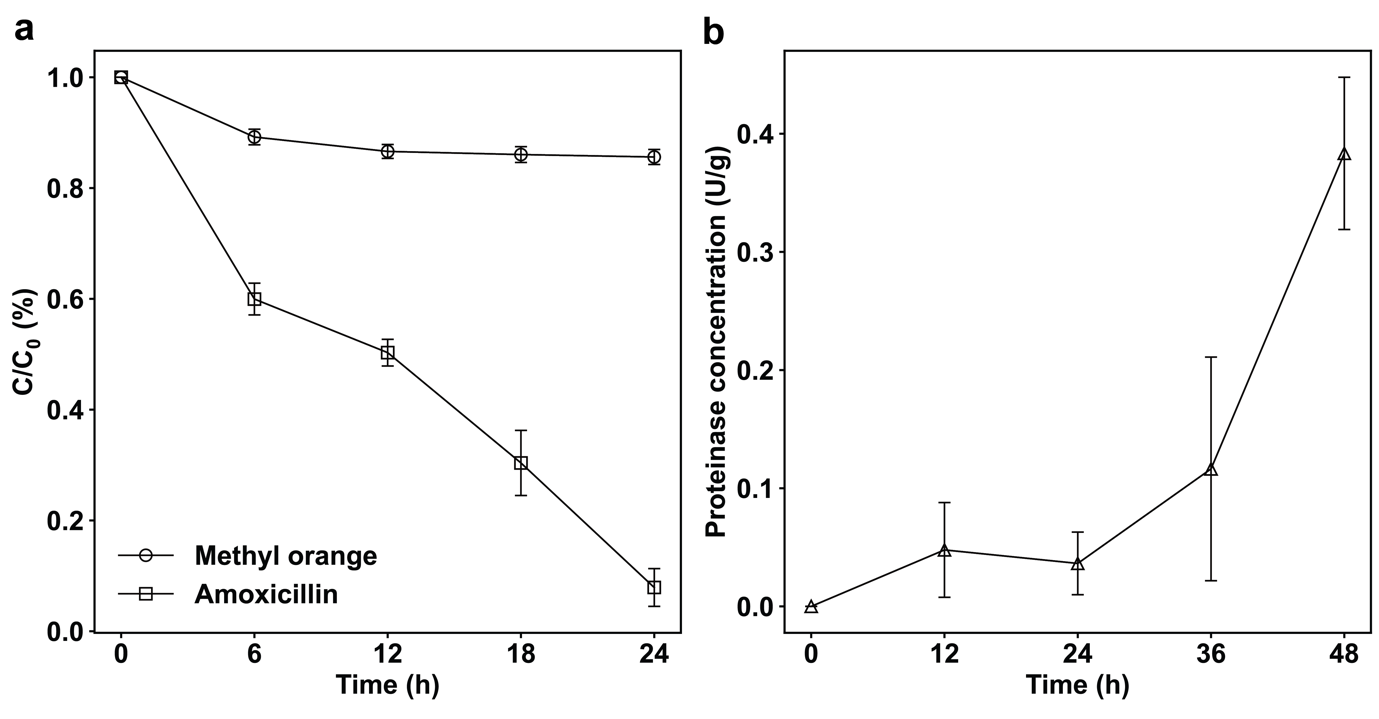

**Fig. 12 the biodegrading capacity of the microalgae-bacteria consortium immobilized in heterogeneous scaffolds.** (a) Removal of methyl orange (100 mg/L) and amoxicillin (300 mg/L) in synthetic wastewater by heterogeneous PAM scaffolds in 24 hours. *n* = 3 independent experiments. (b) The concentration of proteases in the media over time in 48 hours, *n* = 3 independent experiments. All data are presented as mean values +/- standard deviation.

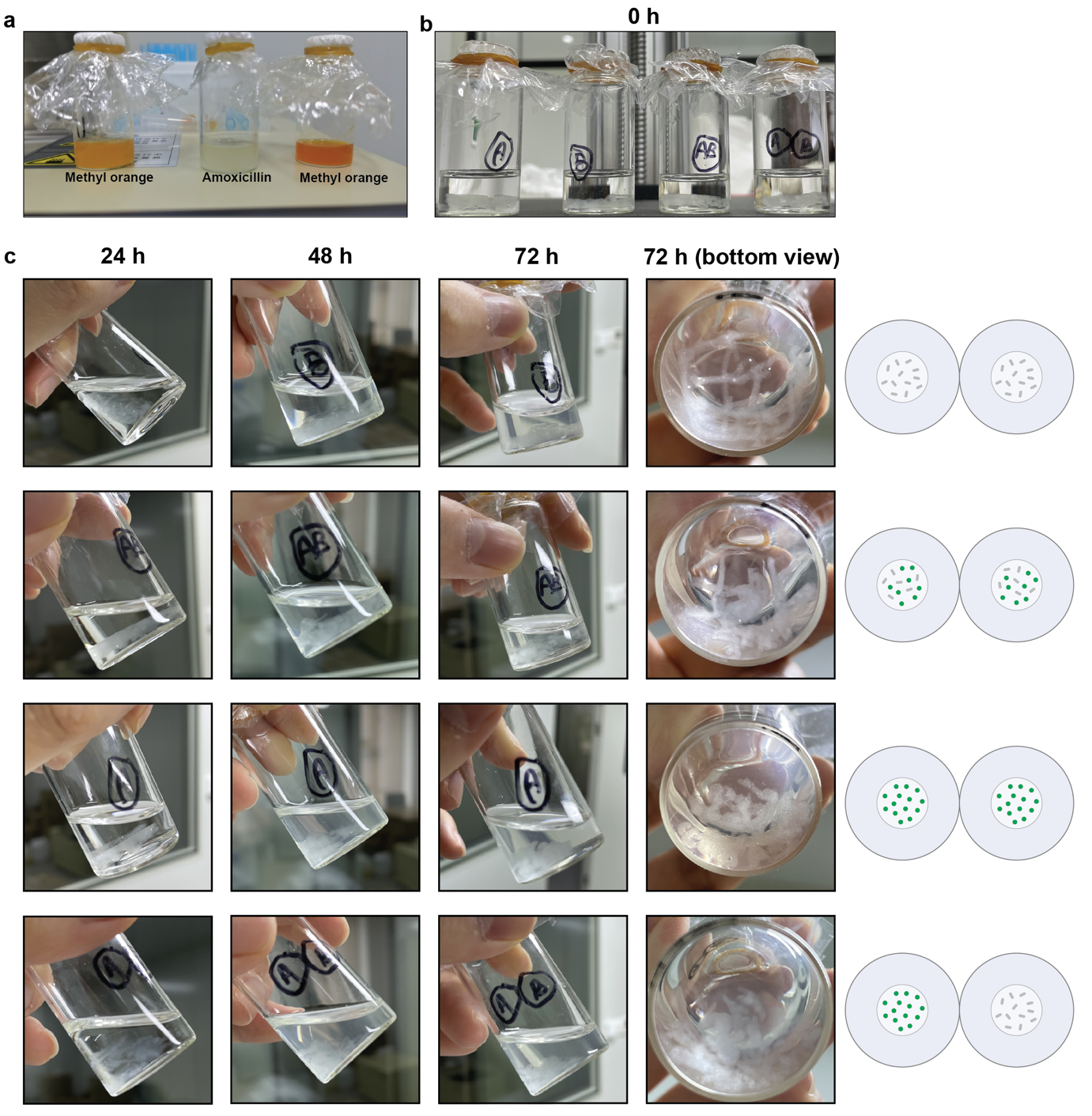

**Fig. 13. Microscopically compartmentalized Microalgae-bacteria co-culture promotes scaffold disintegration.** (a) After 48 hours of incubation, heterogeneous scaffolds immobilizing the microalgae-bacteria in separate microgels were completely liquified, possibly due to proteolytic digestion. (b) Four scaffolds at day 0 before incubation. “A” stands for algae and “B” bacteria. (c) Disintegration of scaffolds. Visual support for Fig. 5e in the main content.

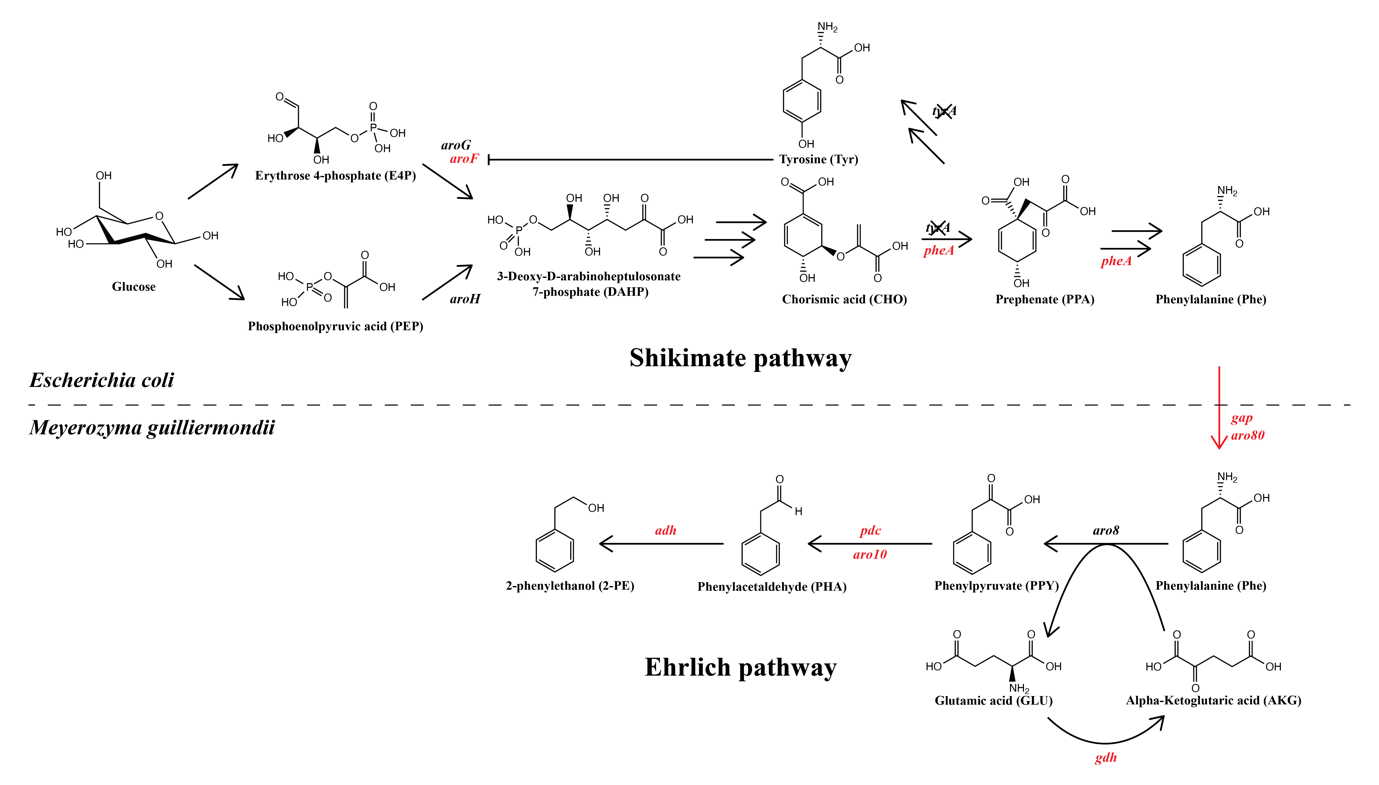

**Fig. 14. Reaction cascade for 2-PE fermentation utilizing metabolically engineered *Escherichia coli-Meyerozyma guilliermondii consortium.*** In light of the intimate connection between the biosynthesis pathway of l-phenylalanine and that of the other aromatic amino acids, especially l-tyrosine, the *pheA* gene, encoding the key enzyme for the biosynthesis of l-phenylalanine, is overexpressed while *the tyrA* gene is knocked out by CRISPR/Cas9, in order to preferentially channel the precursor chorismic acid (CHO) into l-phenylalanine. Additionally, the expression of *aroF*, an essential gene in the shikimate pathway and sensitive to l-tyrosine, is also enhanced^1^. To promote the conversion of l-phenylalanine to 2-PE, genetic modifications are conducted to strengthen the intrinsic Ehrlich pathway of *Meyerozyma guilliermondii*, which comprises three steps: first, l-phenylalanine is transaminated to phenylpyruvate (PPY) with α-ketoglutaric acid (AKG), generating byproduct glutamate (GLU). Accordingly, non-pathway-specific glutamate dehydrogenases (encoded by *the gdh* gene) are overexpressed to increase the supply of AKG. Subsequently, PPY is converted to phenylacetaldehyde (PHA) and further reduced to 2-PE, where key enzymes such as phenylpyruvate decarboxylases (*aro10* and *pdc*) and alcohol dehydrogenases (*adh*) are overexpressed to facilitate the conversion. Besides these genes, *gap*, the general amino acid permeases, and *aro80* that activates the transcription of aromatic amino acid catabolic genes, are also overexpressed to increase the efficiency of bioconversion.

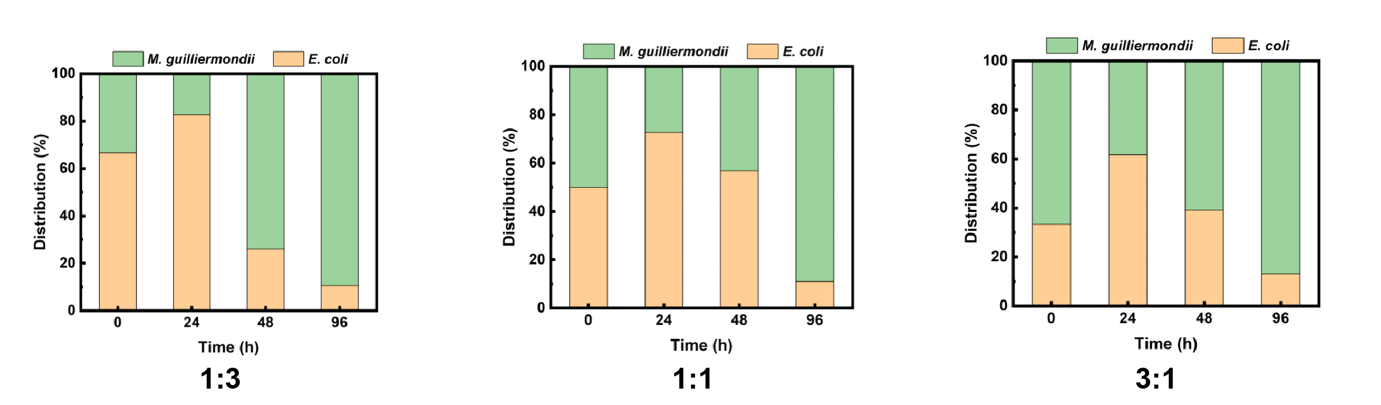

**Fig. 15 Pairwise co-culture of the fungi-bacteria consortium.** The result show that regardless of the initial inoculation ratio, the *M. guilliermondii* becomes dominant in the liquid culture with a ratio of roughly 9:1 between two species in 96 hours.

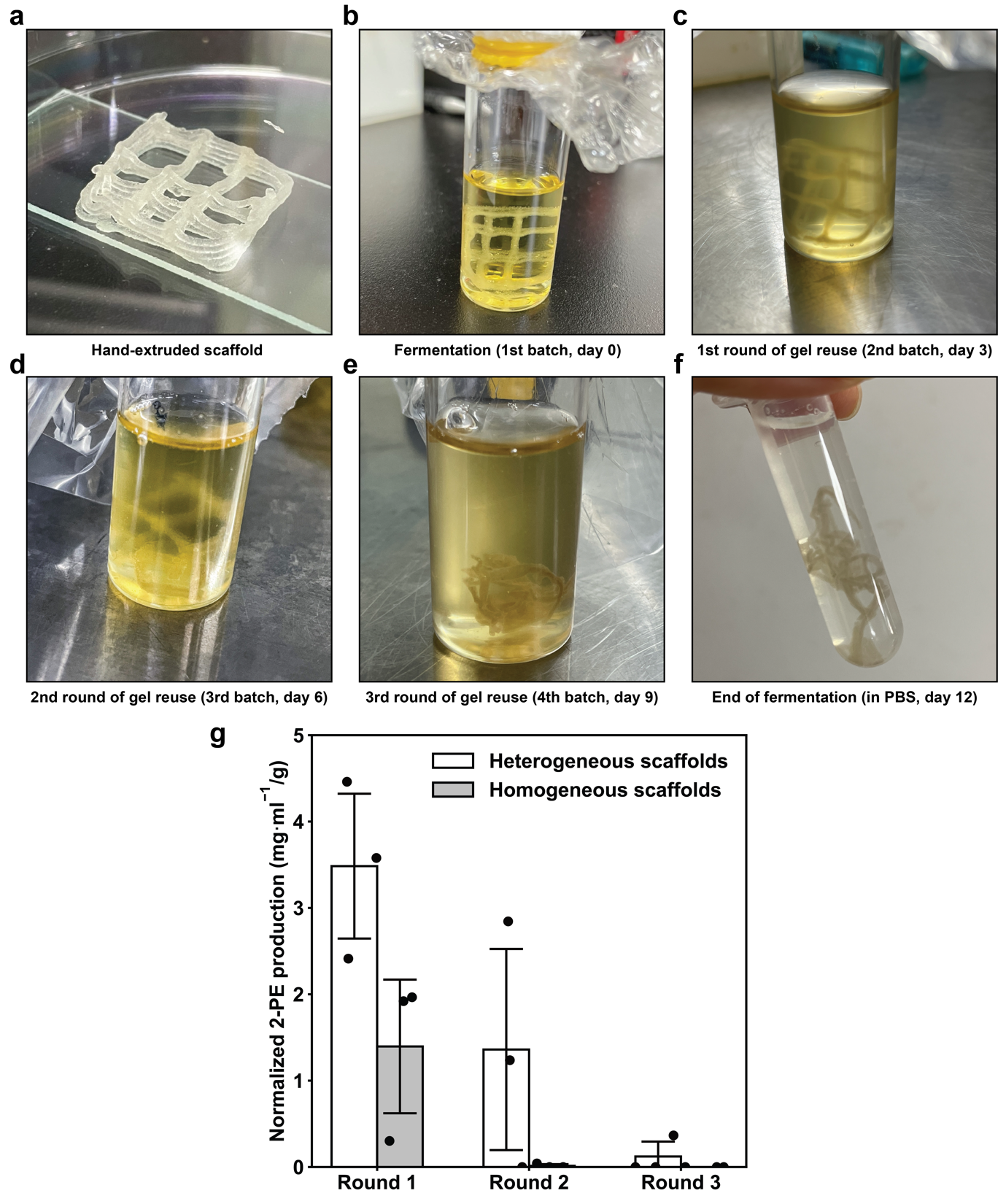

**Fig.16. Reuse of scaffolds for 2-PE batch fermentation.** (a)-(f) Jammed cell-laden core-shell microgels are hand-extruded and patterned into 5-layered 3x3 lattice scaffolds (both heterogeneous and homogeneous) in a biosafety cabinet. Scaffolds are subjected to batch fermentation for a totality of 12 days (four batches, 3 rounds of reuse). Scaffolds start disintegrating after two batches but are not completely comprised at the end of fermentation. (g) Reusability of scaffolds for batch fermentation of 2-PE every 3 days. Data are presented as mean values +/- standard deviation, *n* = 3 independent experiments. After a round of (re)use, scaffolds are washed by fresh media three times before deployed in another batch fermentation.
